# Origin of congenital coronary arterio-ventricular fistulae from anomalous epicardial and myocardial development

**DOI:** 10.1101/2022.01.13.476153

**Authors:** P Palmquist-Gomes, A Ruiz-Villalba, JA Guadix, JP Romero, B Bessiéres, D MacGrogan, L Conejo, A Ortiz, B Picazo, L Houyel, D Gómez-Cabrero, SM Meilhac, JL de la Pompa, JM Pérez-Pomares

## Abstract

**Aims:** In this work we investigated the embryonic origin of coronary arterio-ventricular connections, known as coronary artery fistulas (CAF), a congenital heart disease associated to postnatal and adult changes in systemic hemodynamics that may cause cardiac ischemia.

**Methods and results:** we have used different animal models (mouse and avian embryos) to experimentally model CAF morphogenesis. Conditional *Itga4* (alpha 4 integrin) epicardial deletion in mice and cryocauterisation of chick and quail embryonic hearts disrupted epicardial development and ventricular wall growth, two essential events in coronary embryogenesis. Additional transcriptomics and *in vitro* analyses were performed to better understand how arterio-ventricular connections are originated in the embryonic heart. Our results show myocardial discontinuities in the developing heart of mutant mice presenting epicardial defects and avian embryos submitted to a physical cryodamage of the ventricle. These ventricular discontinuities promote the formation of endocardial pouch-like structures resembling human CAF. The structure of these CAF-like anomalies was compared with histopathological data from a human CAF, showing histomorphological and immunochemical similarities. Both human and mutant mouse hearts showed similar anomalies in the compaction of the ventricular myocardium. In vitro experiments showed the abnormal contact between the epicardium and the endocardium promote the precocious differentiation of epicardial cells to smooth muscle.

**Conclusion:** Our work suggests that myocardial discontinuities in the embryonic ventricular wall are a causative of CAF. These discontinuities would promote the early contact of the endocardium with epicardial-derived coronary progenitors at the cardiac surface, leading to ventricular endocardial extrusion, precocious differentiation of coronary smooth muscle cells, and the formation of pouch-like aberrant coronary-like structures in direct connection with the ventricular lumen.

**Translational perspective:** Congenital coronary artery fistulas (CAFs) lead to complications such as myocardial hypertrophy, endocarditis, heart dilatation and failure. Unfortunately, and despite their clinical relevance, the origins these congenital anomalies remain unknown. In this work, we provide information on the developmental mechanisms involved in the formation of CAFs that is relevant for their early diagnosis and prevention.

## Introduction

Congenital heart disease (CHD) affects around 1% of new-borns^1^, and may have a direct impact on cardiovascular performance during paediatric and adult life^2^. Despite the remarkable social impact of CHD, not much it is known about the embryological defects underlying the origin of many of these conditions. This lack of information on the cellular and molecular mechanisms that cause CHD limits our chances of discovering intrinsic and extrinsic factors that potentially disrupt normal embryonic heart development novel early diagnostic markers for these ailments^3^. We are thus still far from having enough information to design and implement an effective CHD prevention strategy. In this work, we provide experimental, mechanistic evidence on the origin of a distinct form of CHD that affects coronary blood vessels and other cardiac structures.

Coronary artery (CA) anomalies are a well-characterized subset of CHD^4^. Some CA anomalies can severely disrupt cardiac function and even result in sudden cardiac death. It has been reported that 0.6-1.5% of patients undergoing invasive cardiovascular imaging have coronary anomalies, including coronary artery fistulae (CAFs)^4–6^. CAFs are abnormal communications between a coronary artery and other cardiovascular cavity that often are asymptomatic but can also cause long-term variations in the systemic hemodynamics due to the arrest of part of the blood circulation that may result in ischemia. The two main sub-types of CAFs described in the literature involve the connection of a coronary artery to either the lumen of a cardiac chamber (coronary-cameral fistula) or a large cardiac vessel (coronary arterio-venous fistula). The most common CAF type reported in the literature is the arterio-ventricular one (41%), followed by coronary artery connections to the right atrium (26%), the pulmonary trunk (17%), the coronary sinus (7%), the left atrium (5%) and the left ventricle (less than 3%)^7^. In particular, arterio-ventricular fistulae involving large coronary vessels often manifest as a coronary wall convexity (dilation) forming a conspicuous cameral, pouch-like structure at the cardiac surface. Although studies differ in their conclusions on CAF incidence in the right versus the left ventricle^7,8^, solitary ventricular CAFs were found to be more frequent than multiple microfistulas^9^.

It is well established that the vast majority of CAFs identified at birth have a congenital origin^10^. Importantly, these cardiac congenital defects should not be mistaken for isolated connections existing between blood vessels and /or cardiac cavities that result from surgical interventions, traumatic events or infections (e.g. endocarditis), which are often referred-to as ‘fistulae’ and even ‘pseudoaneurisms’^11–13^. Indeed, congenital CAFs are true anomalies that derive from abnormal embryogenesis and, as such, they frequently associate to other forms of CHD, including Tetralogy of Fallot, atrial and ventricular septal defects, persistent ductus arteriosus, pulmonary atresia and left ventricular non-compaction, among others^8^. Some authors have attempted to relate CAFs with gross genetic abnormalities such as chromosome deletions, e.g. 22q11.2^14^, and conditions involving chromosome number variation like Turner, Klinefelter or Down syndromes^15^, but so far the pathogenesis of CAFs remains largely unknown.

Approaching the mechanistic origin of CAFs requires a detailed understanding of embryonic coronary vascular and ventricular wall formation and maturation. Recent studies have demonstrated that coronary arteries form from a primary endothelial capillary network, including cells from the endocardium (sinus venosus and ventricle) and the epicardial progenitors at the septum transversum/proepicardium (ST/PE)^16–18^. This finding implies that, in order to complete the coronary vascular circuitry, cells from the inner ventricular endocardium (preferentially fated to form coronary arteries and capillaries) and the outer epicardium (comprising endothelial cells that will mostly incorporate to coronary veins), merge by crossing the cardiac chamber myocardium that separates them. Moreover, the epicardium, whose contribution to the forming coronary vasculature (intimal endothelium, medial smooth muscle cells and adventitial fibroblasts) is well-known^18–21^, also acts as an instructive signalling centre that promotes the growth and the maturation of the adjacent myocardium^22^. Therefore, it is very likely that both anomalies in the cells that build the coronary vascular tree and the disruption of myocardium growth and maturation are involved in the origin of CA anomalies. In accordance with this hypothesis, animal models for defective epicardial development display a thin compact ventricular myocardium^23^ and coronary defects^24,25^, some of which are reminiscent of CAFs^26^.

In this work, we hypothesize that the disruption of epicardial embryonic development leads to the formation of CAF in mammals by altering embryonic myocardial growth and the differentiation of epicardial derivatives. In order to test this hypothesis, we have conditionally deleted the *Itga4* gene (*α4-integrin*) in epicardial progenitors. This gene encodes for the α4 integrin subunit, whose cardiac expression is restricted to the epicardium. Integrin dimers including α4 integrin subunit promote epicardial adhesion to the myocardium by interacting with the myocardially-expressed Vcam-1 ligand. Since mice with systemic *Itga4* loss-of-function show an extreme cardiac phenotype and die around embryonic day (ED) 9.5-10^27^, we have used the *Cre/LoxP* system to circumvent the early embryonic lethality while preserving the anomalous epicardial formation and the myocardial defects that are secondary to epicardial developmental anomalies. Transcriptomic analysis of mutant mice and micromanipulation of chick embryos have been used to uncover the cellular and molecular mechanisms underlying CAF pathogenesis, and results from these studies were compared with histopathological data from a human paediatric CAF to confirm our experimental findings.

## Methods

All animals used in our research program were handled in compliance with the European and Spanish *guidelines for animal care and* welfare.

### Mice

To circumvent the early lethality of *Itga4* systemic ablation^27,28^, this gene was conditionally deleted in the ST/PE by using a *G2-Gata4-Cre* mouse transgenic line^18^. *Itga4*^flox^ and *G2-Gata4*-Cre mice have been previously described^29,30^. To track epicardial derivatives, *G2-Gata4*-Cre mice were also crossed with *Rosa26R*-YFP mice^18^. Heterozygote *G2-Gata4*^Cre/+^ were bred with *Itga4*^flox/flox^ mice to generate double heterozygous animals that were used to obtain *G2-Gata4*^Cre/+^*;Itga4*^flox/flox^ animals.

### In situ hybridization (ISH)

E9.5 embryos were fixed, embedded in paraffin and sectioned in sterile conditions. In situ hybridization was performed following standard protocols (see supplementary information). The probe was generated with the forward 5’-ATGGTAACCGTAGCTGTACCT-3’ primer and the reverse 5’-AGTCATCCTTGTTCCCACTTG-3’ primer.

### Trichrome staining of paraffin embedded samples

For routine histological analyses, embryos or embryonic hearts were fixed, embedded in paraffin and sectioned. tissue sections were dewaxed in xylene, hydrated in ethanol series and rinsed in distilled water. Samples were then submitted to Mallory’s trichrome staining (see supplementary information).

### Immunohistochemistry in mouse embryos

For immunohistological analyses, embryos were fixed in a 4:1 methanol:DMSO solution overnight at −20°C, dehydrated in an ethanolic series, embedded in paraffin, and sectioned in a microtome. Immunofluorescence of the tissue was performed following standard methods (see supplementary information). Primary and secondary antibodies used in this study are listed in table 1 and table 2, respectively.

### BrdU tissue incorporation, immunohistochemistry and quantification

A working volume of 420-450μl from an aqueous BrdU (Sigma, B9285) stock solution (10mg/mL) was injected intraperitoneally in pregnant females 30 minutes before embryo extraction. Embryos were fixed in a 4:1 methanol:DMSO solution overnight, embedded in paraffin, and sectioned with a microtome. Sections were dewaxed, rehydrated, treated with 2N HCl (30’ at room temperature) and washed again in 100mM sodium tetraborate (5’). Following steps and image acquisition were performed according to standard protocols (see supplementary information). Proliferating cardiomyocytes were estimated with the IMARIS^®^ software (see supplementary information). A t-test and a post-hoc Lubischew coefficient analyses were used to assess statistical significance.

### RNA sequencing (RNA-Seq)

RNA was isolated from heart ventricles of *G2-Gata4*^+/+^*;Itga4*^flox/+^ control and *G2-Gata4*^Cre/+^*;Itga4*^flox/flox^ mutant E11.5 embryos using the Arcturus Picopure RNA isolation kit (Applied Biosystems), including a step for removing genomic DNA. For each genotype, RNA coming from 3 ventricles was pooled (n=4-5). Quality control was performed on an Agilent 2100 bioanalyser using an RNA 6000 Pico LabChip kit (Agilent Technologies). The RNA integrity number (RIN) was 10 for all the samples. cDNA was prepared using the standard Illumina TrueSeq RNASeq library preparation kit. Libraries were sequenced in a GAIIx Illumina sequencer using a 75bp single end elongation protocol. Sequencing read quality was assessed with FastQC (S. Andrews, Babraham Institute). Contaminating Illumina adapters were trimmed with Cutadapt 1.7.1 which also discarded reads that were shorter than 30 bp. The resulting reads were mapped against the mouse transcriptome (GRCm38, release 76) and quantified using RSEM v1.2.20.

Prior to statistical analysis, genes with sufficiently large counts were identified and retained with the function filterByExpr, implemented in the R package edgeR. Data normalization and statistical testing was performed with DESeq2. Enrichment analyses for signalling pathways and gene ontology categories were performed with the R package clusterProfiler. KEGG was selected as the database for pathways and we focused on biological process ontology.

### qPCR validation and data analysis

RNA isolation and qPCR analyses were performed following routine protocols (see supplementary information). The melting curve analysis (ViiA TM 7 Real-Time PCR system, Applied Biosystems) and size fractionation by agarose gel electrophoresis were used to confirm amplification of the expected products. Target quantity (N0) was obtained from the extracted raw data using LinRegPCR program. *Pgk1* and *Ppia* were selected as reference genes as previously described31. Primer sequences (supplementary Table 1) were designed using primer3, BLAST (NIH) and oligo analyser (IDT) software. Graphs and statistical analysis were performed by a non-parametric one-way analysis of variance with a Kruskal-Wallis post-hoc test in GraphPad Prim version 6.0 (GraphPad Software) (P<0.05).

### Protein-protein interaction models

A protein-protein interaction (PPI) sub-network was identified by selecting genes derived from the RNA-seq data analysis, then identifying the predicted proteins associated with these genes and selecting genes and interactions within the first two levels of interaction (A-B-C), considering up to 10 contacts per level (see Supplementary information for further details). As a result, two different sub-networks were identified, one from *VCAM1* and the second from *ITGA4*, representing two different cellular types, cardiomyocytes and epicardial cells, respectively. Interactions between proteins were downloaded from StringDB and visualized in Cytoscape. Nodes were selected based on first and second PPI associations with *VCAM1* and *ITGA4* using a maximum of 10 connections derived from "experiments" or "databases".

### Cryoinjury of the avian embryonic heart

The second experimental model used in this work was the cryocauterisation of the avian embryonic heart. Chick and quail embryos were staged according to the Hamburger and Hamilton stages of chick development^32^ and the classification for quail developmental stages^33^, respectively. Eggs were kept in a rocking incubator at 38°C for 55-60 hours (HH17). A N_2_-cooled copper probe was used to burn the myocardial wall as previously described34. Routine histological analyses were performed as previously explained in mouse embryos (Mallory’s trichrome staining).

### 3D reconstructions of chick damaged hearts

To acquire 3D images, chick hearts were washed in PBS and fixed in paraformaldehyde for several weeks at 4°C. Then, hearts were washed in methanol series, embedded in JB4 resin, sectioned and analysed by a high resolution episcopic microscope (HREM). Two different stages (4 days post-injury “dpi” and 16 dpi) were analysed. The 3D shape of the tissue was reconstructed using the ImageJ^®^ software.

### Immunohistochemistry in avian embryos

Chick and quail embryos were fixed in a 4:1 methanol:DMSO solution overnight, dehydrated in an ethanolic series, embedded in paraffin and sectioned in a microtome. Immunostaining of avian tissues was performed following the same protocol described in the mouse (see supplementary information). For QCPN immunostaining, epitopes were unmasked using TEG buffer (0.12% Trizma base and 0.02% EGTA diluted in distiller water; pH 9) in a pressure cooker for 10 minutes, and the signal was amplified with TSA biotin system kit (PerkinElmer, NEL700A001KT).

### Proepicardium-endocardium co-cultures

Sharp tungsten needles, small iridectomy forceps and scissors were used to isolate quail tubular hearts (HH 16-17) under a dissecting scope. Isolated ventricles and proepicardia were cultured in different conditions (see supplementary data). All samples were fixed overnight in Methanol-DMSO (4:1) solution. Fixed tissue constructs were dehydrated in an ethanol series, washed in butanol, embedded in paraffin and sectioned in a microtome. Some aggregates were not embedded in paraffin and processed as whole mount samples.

For immunohistochemistry, sections were stained following the same protocol than previously described for mouse samples (see supplementary information).

### Necropsy and histological inspection of a human paediatric CAF

The autopsy of a six weeks-old patient with a diagnosis of complex congenital heart disease, who had previously undergone a palliative procedure (Blalock-Taussig shunt) and had died of sudden cardiac arrest, was performed at the Necker Hospital (Paris, France). Adequate informed consent has been obtained for the study of paediatric tissue samples. Post-mortem analysis revealed massive cardiac tamponade as a probable cause of death. Formalin-fixed samples were embedded in paraffin, sectioned with a microtome and mounted on slides. Tissue sections were dewaxed with xylene and hydrated in an ethanolic series. Routine histological analyses were performed by Mallory’s trichrome staining as detailed above. Immunochemical analyses of the human tissue were performed as described in the supplementary information.

## Results

### G2-Gata4^Cre^;Itga4^flox^ mutant mice display epicardial and myocardial developmental defects

To evaluate the role of the epicardium in the origin of CAF, the *G2-Gata4Cre* epicardial driver mouse line, displaying a strong Cre expression in ED 9.5 epicardial progenitor (proepicardial) cells (Fig.1A, B), was crossed with mice carrying floxed *Itga4* alleles (*Itga4*). Crossing these two transgenic lines resulted in the conditional deletion of the *Itga4* gene in epicardial progenitors (*G2-Gata4^Cre/+^*;*Itga4^flox/flox^*, from here onwards, *G2-Itga4* mutants), as shown by the reduced *Itga4* mRNA levels found in *G2-Itga4* mutant proepicardia (Fig.1C, D). E10.5 *G2-Itga4* mutant embryos did not show any evident myocardial wall anomaly, but epicardial formation, highlighted by the presence of cytokeratin-positive cells over the heart surface, was delayed in these animals as compared to stage-matched control embryos (Fig.S1A,B). Epicardial developmental arrest continued to be evident in E11.5 (Fig.1E-H) *G2-Itga4* embryos. At this stage, the compact ventricular myocardium remained abnormally thin in mutant embryos, which displayed discontinuities in their ventricular walls (Fig.1G-H). In accordance with the low numbers of epicardial cells found in *G2-Itga4* mutant embryos, the expression of *Itga4* and epicardial-related genes like the *Wilms’ tumour suppressor 1* (*Wt1*) and the transcription factor *Tcf21* was sharply reduced in mutant ventricles as compared with control ones (Fig.1I-L). This E11.5 epicardial growth arrest was evidenced by the absence of Wt1 protein on the heart surface (Fig.S1C,D). At ED12.5 *G2-Itga4* mutant embryos often presented evident pericardial haemorrhage, as shown by gross anatomical and histological examinations (Fig.S1E-G). The ventricular myocardial discontinuities found in the ventricles allowed for the local extrusion of the endocardium (Endomucin staining) towards the pericardial cavity forming characteristic pouch-like structures (Fig.1M, N). Immunostaining confirmed the efficient deletion of *Itga4* in the epicardium of mutant embryos (Fig.S1H-I’). At E13.5, all mutant embryos retrieved were dead.

**Figure 1.**
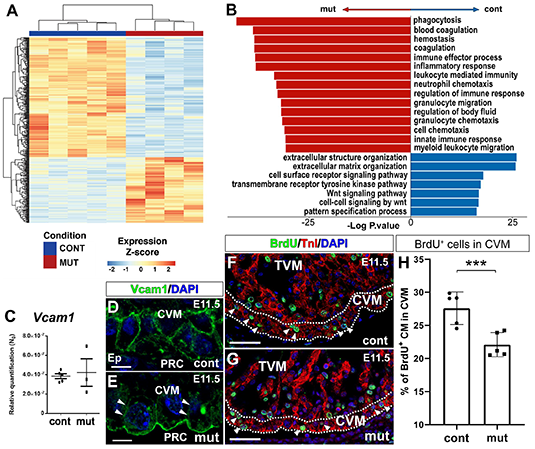
*Itga4* (α-4 integrin) disrupts epicardial and myocardial embryonic development and results in endocardial extrusions. **A, B**. YFP reporter expression in *G2-Gata4*^Cre^*;Rosa26R*-YFP mice highlights epicardial progenitor cells at the septum transversum/proepicardium (green fluorescence, arrow in A, boxed area in B-B’). **C, D**. *G2Gata4CRE*-mediated *Itga4* deletion in the ST/PE severely reduces *Itga4* mRNA expression in mutant (*G2-Gata4*^Cre/+^;*Itga4*^flox/flox^) proepicardium (compare the boxed areas in C,C’ and D,D’). **E-H**. E11.5 mutant mice display impaired epicardial (Ck^+^, green fluorescence) development (compare E-F with G-H). Ct3 counterstains the myocardium. Ventricular myocardial discontinuities are frequently found in mutant embryos (H, arrowheads). **I-L**. RT-qPCR shows a significant (P<0.05) loss of *Itga4* gene expression in E11.5 mutant ventricles (mut; n=4) when compared to control ones (n=5 in I; n=4 in J), (I,J). The expression of the epicardial marker genes *Wt1* (K) and *Tcf21* (K, L, respectively) is also sharply reduced in mutant ventricles (n=4) as compared to control ones (n=5) (P<0.05). Each replicate represents a pool of three ventricles. Statistical significance was obtained by a non-parametric one-way analysis of variance with a Kruskal-Wallis post-hoc test. **M, N**. At E12.5, the mutant ventricular endocardium (End^+^) locally reaches the pericardial cavity and forms vesicular structures (asterisks indicate the inner lumen, compare M,M’ with N,N’). The dashed line in N,N’ marks the endocardial extrusion path through a myocardial discontinuity. The ventricular lumen is in communication with the lumen of the endocardial vesicle. **Abbreviations**: Ck, cytokeratin; Ct3, cardiac troponin T; CVM, compact ventricular myocardium; DAPI, diamidino-2-phenylindole; End, endomucin; ENDO, endocardium; EP, epicardium; LV, left ventricle; PE, proepicardium; PRC, pericardial cavity; RV, right ventricle; ST/PE, Septum transversum/proepicardium; TnI, troponin I; TVM, trabeculated ventricular myocardium; V, ventricle; YFP, yellow fluorescent protein. **Scale bars:**A,B,C,D,E,G: 100μm; B’,C’,D’,E’,F,G’,H,M,N: 50μm; M’,N’: 25μm.

### Epicardial *Itga4* deletion alters the transcriptome of developing cardiac ventricles

To identify the altered signalling pathways underlying the ventricular defects present in *G2-Itga4* mutants, we performed RNA-sequencing on microdissected ventricles from E11.5 controls (*G2-Gata4*^+/+^;*G2-Itga4*^flox/+^) and mutant (*G2-Itga4*) embryos. A total of 376 differentially expressed genes (DEG) (p.value<1e^−3^) were identified after comparing mutant versus control samples. Out of those 376 genes, 134 were overexpressed and 242 underexpressed in the mutant tissues (Fig.2A and Fig.S2). Gene Ontology (GO) enrichment analysis revealed functional differences between the two genotypes (Suppl. excel file 1). Specific extracellular matrix (ECM) related genes (*Col1a1*, *Col1a2*, *Col8a1*, *Col8a2* or *Itga8*), which are annotated to GO terms such as ‘*Extracellular structure organization*’ and ‘*Extracellular matrix organization*’, were downregulated in mutant ventricles. However, genes annotated in GO categories as ‘*Inflammatory response*’ (*Ccl6*, *Cd68*), ‘*Immune Effector Process’ (ItgaI, Vav1), ‘Blood coagulation*’ (*F2rl2*, *Nbeal2*) or ‘*Wound healing*’ (*Ptpn6*, *Trim72*) were upregulated. The pathway enrichment analysis also revealed that ‘*Focal adhesion*’ and ‘*ECM-receptor interaction*’ were the most significantly enriched pathways in control animals (Fig.2B). These data suggested that the abrogation of epicardial-myocardial cell-to-cell and cell-to-matrix interaction could, at least, partially explain the ventricular phenotype of mutants (see Fig.1). Prompted by this finding, we investigated the status of *Vcam1* gene, encoding for the canonical myocardial receptor of α4-integrin. While no significant differences were found in the expression of *Vcam1* gene between mutant and control ventricles (Fig.2C), the altered distribution of Vcam1 in the plasma membrane of mutant cardiomyocytes was evident (E11.5) (Fig.2D,E). This finding indicates that anomalous Vcam1-positive cells distribution in mutant cardiomyocytes did not result from altered *Vcam1* transcription, but rather derives from a larger scale change in cardiomyocyte adhesion to the ECM or to other cells. To identify potential molecular mechanisms underlying the role of the Vcam1/α4-integrin axis in the mutant ventricular phenotype, a protein-protein interaction network model analysis was conducted (see Methods). The analysis identified two sub-networks of protein interactions (Ezrin(Ezr)/Radixin/Moesin(Msn) and Ncf/Cybb/Rac1) that could be altered in *G2-Itga4* embryos (Fig.S3A). Each sub-network contains proteins that potentially interact with *ITGA4 via VCAM1* (Fig.S3B), and may thus affect the polarization of the cardiomyocytes as consequence of the absence of epicardial ITGA4 or altered cell adhesion^35^.

**Figure 2.**
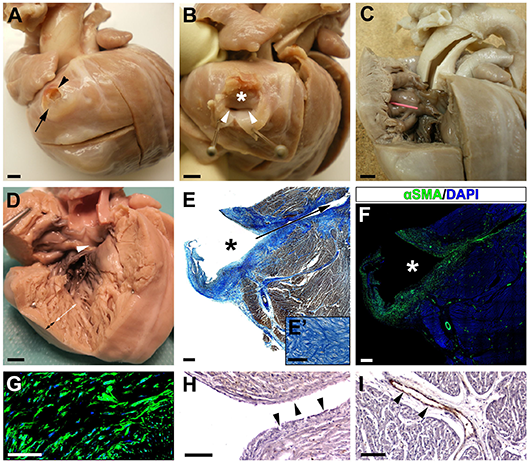
*Itga4* epicardial deletion modifies the transcriptomic and structural profile of embryonic ventricles. **A**. Comparative RNA-seq analysis shows that 376 genes are differentially expressed (DEG) between mutant (*G2Gata4*^CRE/+^;*Itga4*^flox/flox^) (n=4) and control (*G2Gata4*^+/+^;*Itga4*^flox/*^) (n=5) E11.5 ventricles (p.value < 1e^−3^); 134 genes are increased and 242 are decreased. each replicate represents a pool of three ventricles. **B**. Gene Ontology (GO) enrichment analysis (an Over Representation Analysis with a Hypergeometric distibution with a Benjamini-Hochberg adjustment for multiple comparison) reveals important differences between the two genotypes. Pathway enrichment is expressed as the –log [P] adjusted for multiple comparison. The direction of the bars indicates which category was over/underrepresented. All categories in red are overrepresented in mutant animal, whereas all categories in blue are overrepresented in control animals. **C-E**. RT-qPCR analysis of E11.5 embryonic ventricles (n=5 in control and n=3 in mutant; each replicate represents a pool of three ventricles) does not reveal significant variations in *Vcam1* gene expression (C), but Vcam1 protein mislocalisation is evident in ventricular cardiomyocytes (compare D with E; arrowheads mark the reduced Vcam1 distribution in the lateral plasma membrane of cardiomyocytes). Statistical significance was obtained by a non-parametric one-way analysis of variance with a Kruskal-Wallis post-hoc test (P<0.05). **F-H**. BrdU uptake (F,G, arrowheads) is significantly reduced in E11.5 mutant ventricular compact myocardium (dashed lines) (n=5) as compared with stage-matched control (n=5) (P<0.01, 3 asterisks, H). The statistical significance was assessed by a t-test and a post-hoc Lubischew coefficient analysis. For the quantification of BrdU-positive cells, each replicate represents the mean value from quantifications in three ventricular sections. **Abbreviations**: BrdU, bromodeoxyuridine; CVM, compact ventricular myocardium; DAPI, diamidino-2-phenylindole; TnI, troponin I; TVM, trabeculated ventricular myocardium. **Scale bars**: D,E: 25μm; F,G: 50μm.

Finally, to evaluate whether epicardial depletion in mutant hearts affected cardiomyocyte proliferation, BrdU experiments were carried out. BrdU uptake was studied in the compact ventricular myocardium of mutant embryos at E11.5 (Fig.2F,G). This analysis revealed a significant (p<0.01) decrease in cycling cardiomyocytes of the compact ventricular layer (Fig.2H).

### Histopathology of a paediatric human CAF

The autopsy of a six weeks-old patient who died of sudden cardiac arrest at home revealed the presence of several structural cardiac congenital anomalies, including double outlet left ventricle (DOLV) with outlet ventricular septal defect (VSD), and a large coronary arterio-ventricular fistula with a pouch-like appearance (Fig.3A,B). This CAF was found at the cardiac surface, near the left anterior descending coronary artery. The outer wall of the CAF displayed a patent rupture (Fig.3A,B), and its inner cavity was directly connected to the right ventricular lumen through two independent orifices (Fig.3B, C). The right ventricle was poorly compacted, displaying large trabecular processes (Fig.3D). The histological analysis of the CAF wall using Mallory’s trichrome staining showed the accumulation of ECM proteins in the wall of the structure (Fig.3E). ECM accumulation overlapped with the presence of large, compacted numbers of smooth muscle cells in the CAF wall and the medial layer of associated coronary arteries (Fig.3F,G). To determine whether the CAF was lined internally by endothelial cells the expression of Von Willebrand (VW) factor was considered. Anti-VW immunohistochemistry indicated little or no immunoreactivity for this marker in the tissue covering the CAF lumen (Fig.3H), whereas significant amounts of VW protein were detected in the endothelium of adjacent coronary vessels (Fig.3I).

**Figure 3.**
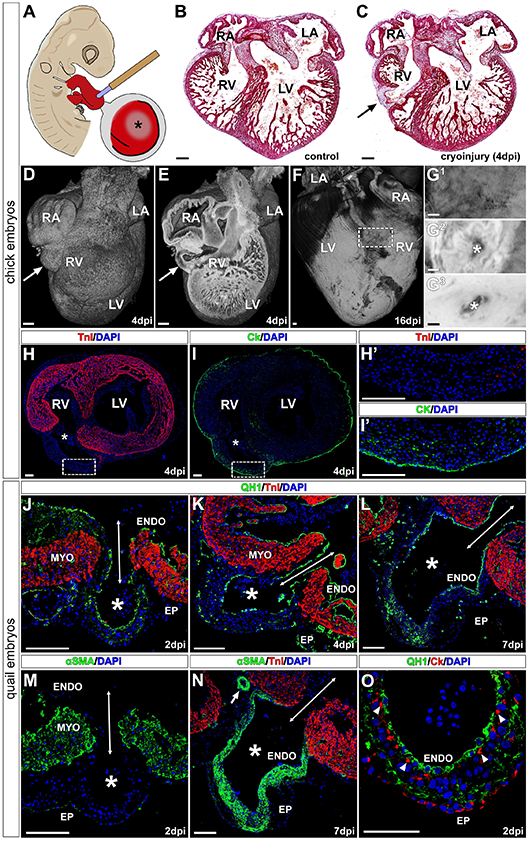
Necropsy of a paediatric CAF case. **A-C.**Right lateral view of a six weeks-old human heart showing a coronary arterio-venticular communication or fistula (CAF) with a characteristic pouch-like appearance (A, arrow). The wall of the fistula was found to be ruptured at the time of necropsy (A, arrowhead). The cavity of the fistula (B, asterisk) shows two orifices (B, arrowheads) that penetrate the ventricular wall (B, arrowheads) and are in continuity with the right ventricular lumen (C, pink probe). **D**. Evidence of a large outlet ventricular septal defect (D, arrowhead), and a non-compacted right ventricular apex (D, the white double-headed arrow marks the non-compacted myocardium and the black one the compacted myocardium). **E-G**. Mallory’s trichrome staining of histological sections from this CAF (the asterisk marks the lumen of the fistula and the arrow the communication with the ventricular lumen) shows the accumulation of ECM (E’) in the wall of the structure. Anti-αSMA^+^ cells are very abundant in the fistular wall (F-G). **H-I**. Von Willebrand factor is almost undetectable in the endothelial lining of the CAF (H, arrowheads), but it is conspicuous in the endothelium of the surrounding coronary vessels (I, brown, arrowheads). Nuclei were counterstained with DAPI (F,G; blue). **Scale bars:**A,B,C,D: 1mm; E,F: 500μm; E’, G,H,I: 100μm.

### Cryoinjury of the avian embryonic heart disrupts ventricular myocardial continuity and results in CAF-like structures

Since *G2-Itga4* mouse mutant embryos died shortly after endocardial vesicles form on the heart surface (see Fig.1N), a cardiac cryoinjury method was applied to chick embryonic hearts to better understand the cellular mechanisms underlying the formation of the fistular wall (Fig. 4A). The local cryoinjury of the embryonic ventricular myocardium resulted in a single myocardial discontinuity and the formation of CAF-like structures on the ventricular surface (Fig.4B,C and Fig.S4A-C). A 3D analysis of chick cryodamaged hearts at 4 and 16 days post injury (dpi) (corresponding to HH46 stage, 1 day before hatching), revealed the prevalence of a thin channel (fistula) connecting the pouch-like structure with the ventricular lumen (Fig.4D-G). Immunohistochemical analysis of these structures showed they are limited by an external wall including Cytokeratin^+^ epicardial epithelial cells and epicardial-derived mesenchyme (Fig.4H-I′).

**Figure 4.**
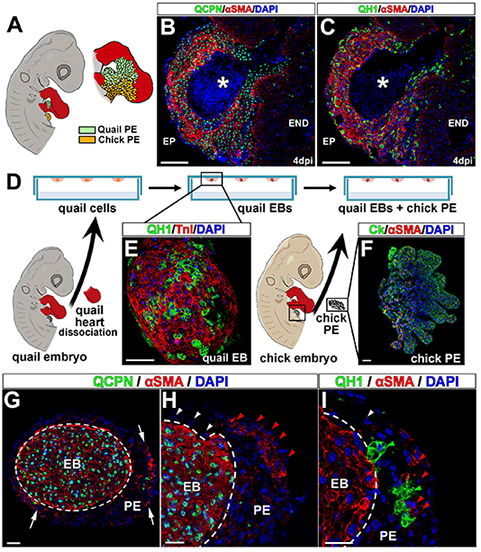
Local cardiomyocyte damage disrupts ventricular myocardial continuity and results in endocardial extrusion towards the pericardial cavity. **A.**Chick embryos were submitted to ventricular cryoinjury at HH16-17 stages. **B,C**. Frontal (cranial) sections of control (B) and cryodamaged (C) chick. The damaged area in the cryoinjured ventricular wall (4 days post injury) is marked with an arrow. **D-G^3^**. HREM reconstructions of cryoinjured chick embryonic hearts. At 4 days post injury (dpi), a pouch-like structure is formed in the damaged area (D,E, arrows) directly connected to the ventricle lumen (E, dashed line). These kind of structures remain visible at 16 dpi (F,G). Transmural analysis of the ventricle shows that the lumen of this fistular vesicle is in communication with the ventricular one (G^1^-G^3^, asterisks). **H,I**. Myocardial ventricular discontinuities (asterisks) are TnI^−^(H,H’) but are externally lined by Ck^+^ tissue (I,I’). **J-O**. Cryocauterisation of quail hearts results in the formation of pouch-like structures also form at the damage site (J-O, asterisks); their lumen is continuous with the ventricular one (J-N, double headed arrows). The quail specific marker QH1 labels the embryonic endocardium and vascular endothelium (J-L, O). At 2dpi, alpha smooth muscle actin (αSMA) expressed in the immature ventricular myocardium (M). αSMA is mainly observed in the wall of fistula-like structures at 7 dpi (N) and the tunica media of adjacent coronary arteries (N, arrow). Endocardial (QH1^+^)-epicardial (Ck^+^) contact takes place early in the damaged area (2 dpi, O, arrowheads). **Abbreviations**: αSMA, alpha smooth muscle actin; CK, cytokeratin; DAPI, diamidino-2-phenylindole; dpi, days post injury; ENDO, endocardium; EP, epicardium; LA, Left atrium; LV, left ventricle; MYO, myocardium; RA, Right atrium; RV, right ventricle; TNI, troponin I. **Scale bars:**B,C,D,E,F,G: 200μm; H,I,H’,I’,J,K,L,M,N,O: 100μm.

In order to characterize the CAF-like endothelial lining, the heart cryoinjury method was also applied to quail embryos, and the QH1 quail endothelial marker was used to counterstain endothelial cells (including endocardial ones) in the developing heart (Fig.4J-L). Three developmental stages of quail embryos were analysed, namely 2-, 4- and 7-dpi (Fig.4J-O). As described for chick embryos (a species for which no *bona fide* endothelial markers are available), the continuity of the myocardium was locally lost in cryoinjured quail embryos as illustrated by the absence of troponin (TNI) (Fig.4J) and alpha smooth muscle actin (αSMA) staining in the damaged area (Fig.4M). A local extrusion of putative endocardial and epicardial tissues towards the pericardial cavity was observed in all cryodamaged hearts (Fig.4J-O). Although at 2 dpi αSMA expression can still be found in the embryonic myocardium (Fig.4M), αSMA cardiac becomes progressively restricted to the smooth muscle cells of the developing coronary arteries and the forming wall of CAF-like structures at 4 and 7 dpi (Fig.4N,O). At 2dpi, the extruded endocardium was in contact with some epicardial cells (CK^+^) (Fig.4Q).

### Experimental avian fistulas originate from epicardial-derived and endocardial cells

Interspecific quail-to-chick transplantations have been for long regarded as a reliable method to permanently trace the fate of embryonic tissues^19^ (Fig.5A). Hence, we used this approach to evaluate whether CAF share, at least in part, an epicardial origin with coronary blood vessels^18^. Proepicardial (PE) quail transplantations into chick hosts and subsequent cryocauterisation of the developing chimeric embryo revealed that the majority of smooth muscle cells (αSMA+) forming the CAF-like wall were donor-derived (quail) cells expressing the quail-specific QCPN nuclear marker (Fig.5B). The absence of this QCPN marker (Fig.5B) or the quail-specific endothelial marker QH1 (Fig.5C) in the inner lining of CAF-like structures (which is continuous with the host chick endocardium), suggests an endocardial origin for these cells and rules out PE endothelial contribution to these anomalies.

**Figure 5.**
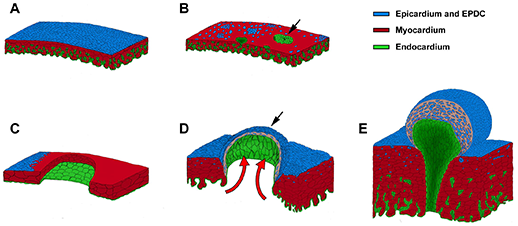
Cellular origins in avian CAF-like structures. **A.**Quail-to-chick proepicardial chimeras were first constructed by grafting a quail proepicardium close to a chick cryodamaged heart and then submitted to cryocauterisation as shown (Fig.4A). All donor (quail) proepicardial-derived cells are QCPN^+^ (B, green fluorescent nuclei) and quail-derived endocardial/endothelial cells are QH1^+^ (C, green fluorescent cells). **D-G**. Co-culture of quail endocardial and chick epicardial tissues. Quail heart dissociation and subsequent hanging drop culture allows for the externalization of quail endothelial cells on the surface of forming embryonic bodies (E, QH1^+^, green fluorescent cells). Co-culture of these embryonic bodies with freshly excised epicardial progenitor (proepicardial) cells (F, Ck^+^/αSMA^−^, green fluorescent cells) results in the rapid differentiation of αSMA^+^ cells from this latter tissue (g, arrows). **H-I**. Chick proepicardial cells (QCPN^−^, blue fluorescent nuclei) cover quail-derived tissues after 1 day of culture (H, dashed line). These proepicardial-derived cells mostly are αSMA^−^(H’, white arrowheads), but some of them are αSMA^+^ (H, red arrowheads). These αSMA^+^ cells (I, red arrowheads) are adjacent to quail endothelial (QH1^+^) cells (I, green arrowheads). Quail immature myocardium is always αSMA^+^/QCPN^+^ (G, H, green fluorescent cell nuclei, red fluorescent cytoplasm). **Abbreviations**: αSMA, alpha smooth muscle actin; Ck, cytokeratin; DAPI, diamidino-2-phenylindole; EB, embryonic body; PE, proepicardium. **Scale bars:**B,C: 100μm; E,F,G,H,I: 25μm.

Finally, to test whether early epicardial cell contact with the embryonic endocardium accelerates smooth muscle differentiation contributing to CAF-like wall formation, interspecific co-culture analyses of chick epicardial progenitor (proepicardium) and quail myocardial-endocardial tissues were developed. Hanging drop culture of quail embryonic hearts (Fig.5D) allowed for endocardial (QH1^+^) growth and accumulation at the surface of the cardiac explant (Fig.5E). Chimeric constructs were formed by aggregating chick proepicardia (Fig.5F) to these quail explants for 24 hours (Fig.5G-I). After immunohistochemical inspection, the alpha-smooth muscle actin (αSMA) marker was observed in chick (QCPN^−^) proepicardial-derived cells in close association with quail endocardial (QH1^+^) cells. αSMA marker expression was scarce in areas where chick cells were in contact with myocardial (αSMA^+^/QH1^−^) but not endocardial (αSMA^−^/QH1^+^) cells (Fig.5G-I).

## Discussion

Aberrant connections between coronary arteries and a heart chamber or a large vessel, commonly referred-to as coronary arterio-ventricular fistulae (CAFs) ^6^, have been known for more than a century. These anomalies are rare, but often associate to multiple cardiac complications such as myocardial hypertrophy, endocarditis, heart dilatation and cardiac failure^4,7,8^.

A certain degree of confusion exists in the literature about the origin of CAFs. This is, at least in part, due to semantics, since the terms coronary ‘aneurysm’ and ‘pseudoaneurysm’ are often interchangeably used to refer to CAF^5,36^. This requires a primary distinction between anomalies of the coronary wall with an acquired origin, which may result from defective surgical procedures^37^ and congenital CAF anomalies with a developmental origin^7^. A note on the nature of CAF should also be made as based on strict histopathological criteria, as true congenital vascular aneurysms involve the local bulging of the three tissue layers of a vessel (intima, media and adventitia), whereas pseudoaneurysms (or false aneurysms) are anomalous structures with a characteristic pouch-like appearance containing tissue from the medial and adventitial mural layers only^38^. In our work, we have revisited the histoarchitecture of human CAF through the histological analysis of a conspicuous CAF found in a six-week-old child. This examination indicates that endothelial (intimal), smooth muscle (medial) and fibroblastic (adventitial) tissues form the wall of these structures. Detailed reports indicate that the wall in human CAFs contains smooth (but not striated) muscle bundles frequently separated by several internal elastic laminae, as well as a characteristic intimal layer showing non-specific fibrous thickening^39^. Similar histological features were found in the paediatric case reported in our study: the internal surface of the CAF is lined by a vascular endothelium displaying a characteristically poor Von Willebrand immunoreactivity associated to the endocardium^40^, with the majority of the fistular wall formed by smooth muscle cells (α-SMA^+^). On a final note, our analysis of this human case revealed that the ventricular myocardium was poorly compacted. This finding developmentally associates poor ventricular thickening or compaction^41^, with the occurrence of congenital CAF as described in paediatric and adult human patients^42^.

While the congenital origin of CAFs is widely accepted^43^, no data on the disrupted developmental mechanisms that lead to CAF formation are available in the literature. Although the anatomical description of CAF may suggest these anomalies are secondary to disrupted coronary vasculature morphogenesis^44^, it has also been suggested that they could form by the persistence of sinusoidal connections between the forming coronary arteries and the primitive tubular heart lumen^45^. These two hypotheses, reinterpreted in light of recent discoveries on the embryonic origin of coronary blood vessels^16–18^, are fully compatible and could explain the origin of part of congenital CAF. Indeed, connections between both the embryonic sinus venosus at the venous pole of the heart and the ventricular lumen with the primitive coronary capillary plexus occurs during normal coronary embryonic development^16–18^. Such connections are progressively lost as the ventricles thicken and mature during the perinatal stages^46^. Thus, the proper growth of ventricular myocardium is necessary to maintain the patterning and stabilization of embryonic coronary vessels and is also required to separate coronary endothelium from its endocardial origin. In this work we suggest that the anomalous disruption of compact myocardial wall formation may result in persistent communications between coronary blood vessels and the cardiac lumen leading to CAF formation.

To test our working hypothesis, we used two animal models. The first one is a mouse with conditional epicardial deletion of the *Itga4* gene. This mutant recapitulate the severe ventricular myocardial compact layer and epicardial phenotype of *Itga4* systemic mutant^27^ but survive beyond its early lethality. Relevant to our approach, *Itga4* mutants represent an extreme case of cell (Itga4) and non-cell autonomous (Vcam1) defects associated to defective epicardial development, a key developmental process required for both coronary blood vessels and ventricular wall formation^23^.

Our second animal model is the chicken embryo, which is amenable to a large variety of *in ovo* experimental manipulations with little or no impact on embryo viability^47^. In particular, we have taken advantage of a recently described procedure allowing for the creation of restricted damage on embryonic tissues by means of local cryocauterisation^34^. In this case, cryocauterisation was used to locally disrupt ventricular myocardial wall continuity.

Genetic deletion of the *Itga4* gene in mouse epicardial progenitors and epicardial cells disrupts early epicardial development, significantly reducing the number of epicardial cells as shown by histological and epicardial-specific gene expression analyses. Abnormal epicardial development in these mutants is concomitant with the lack of thickening of the ventricular compact myocardial layer due to reduced cardiomyocyte proliferation, has been described for other epicardial mutants^48^. Such discontinuities allow for the extrusion of the endocardium, which can be distinguished from the forming coronary vascular endothelium by its strong endomucin expression. Indeed, endomucin marker expression confirms the endocardial origin of the tissue lining the CAF cavity, also suggesting it does not derive from the differentiating coronary vascular endothelium found in the cardiac surface (endomucin-negative).

The transcriptomic analysis of epicardial *Itga4* mutants revealed a significant revealed a significant increase in genes associated to tissue damage (thrombosis, blood clot formation, inflammation pathways) as well as a marked decrease in genes linked to cell surface signalling and ECM organization. These findings, taken together, strongly suggest that epicardial-to-myocardial adhesion mechanisms, including those mediated by the ECM, were affected by *Itga4* deletion. Consequently, we checked the status of *Vcam*-1, the canonical α-4 integrin epicardial ligand^49^. Interestingly, *Vcam-1* gene expression level was unchanged in *Itga4* epicardial mutants, but Vcam1 protein distribution in the plasma membrane of cardiomyocytes was found to be severely altered in these animals. Both the reduced number of epicardial differentiating cells and the altered secretion of subepicardial matrix proteins in mutant embryos could explain the poor cell surface organization of Vcam1. Our in silico protein interaction modelling suggested that different proteins belonging both the Ezrin/Radixin/Moesin and Ncf/Cybb/Rac1 pathways could potentially interact with Vcam1. These pathways could be involved in the developmental origin of the mutant phenotype by regulating ventricular cardiomyocyte polarity. The specific cellular context of genes related to *Itga4* is suggested by the high prevalence of epithelial (and epicardial) specific proteins in the model. Moreover, our model suggests the existence of a biochemical/functional relationship between Itga3 and Itga4. Whether anomalies in Itga4-Itga3 protein interactions could promote the previously described CAF-like phenotype needs further experimental analyses. Taken together, these results indicate that defective cell-to-cell and cell-to-matrix adhesion could underlie the appearance of the characteristic ventricular myocardial discontinuities of epicardial *Itga4* mutants. In turn, these ventricular myocardial wall discontinuities could represent a passage through the compact myocardium for the endocardium, providing a physical continuum between the endocardial lining of the cardiac chamber with the pericardial space.

The death of *Itga4* epicardial mouse mutants at E12.5-13 prevented us from studying the formation of the CAF-like structures in these animals in more detail. To progress in our analysis of these anomalies, we decided to use chick embryos, a highly manipulable experimental animal model extensively used to study embryogenesis. Hence, we carried out limited cryocauterisation of the chick embryonic heart (HH16-17, around 60 hours of incubation) to reproduce the ventricular myocardial discontinuities we identified in *Itga4* epicardial mutants. Our results show that the mechanical disruption of the chick embryonic myocardium faithfully reproduces the formation of arterio-ventricular anomalies showing a structure very similar to that described for human CAF. The combination of cryocauterisation and interspecific quail proepicardium-to-chick chimerisation for cell fate tracking demonstrates that the smooth muscle wall covering these anomalies, as that of coronary arteries, has epicardial origin ^23^. Our results also confirm that the inner lining of the fistula cavity is not an epicardial derivative; the absence of the QCPN quail donor marker and the continuity of these cells with the ventricular endocardium strongly suggest a host endocardial origin for this tissue. Since coronary smooth muscle is known to be a major embryonic epicardium derivative^23^, and endothelial-to-mesodermal progenitor cell contact and paracrine molecular cross-talk efficiently result in smooth muscle differentiation^50^, we tested *in vitro* whether early endocardium-to-epicardial-derived cell contact could effectively induce smooth muscle differentiation from the latter cell type. Our results convincingly demonstrate that early *in vitro* endocardium-to-epicardial-derived mesenchymal cell interaction triggers epicardial cells differentiation into smooth muscle cells, as normally happens *in vivo*.

We have herein shown that myocardial discontinuities in the embryonic ventricular wall, either secondary to anomalous epicardial development or resulting from a direct damage to the embryonic myocardium, cause CAF in two different vertebrate models via the extrusion of the endocardial lining of the developing heart, the precocious and anomalous formation of a smooth muscle wall over the extruded endocardium, and the persistence of arterial to ventricular lumen connection (Fig.6). The histological analysis of a human CAF, compared to our cell-tracing experiments, confirm the nature and origin of the tissues that form this congenital CA anomaly. The experimental analysis of coronary and ventricular wall morphogenesis in two classical animal models for the study of cardiovascular development (mouse and chick) reveals significant similarities in the phenotypes resulting from the combined disruption of epicardial and myocardial normal development that, in turn, strikingly reproduce human coronary arterio-ventricular fistulae. To the best of our knowledge, these results are one of the very few examples on the origin of a specific CHD to be found in the literature, providing clues to the identification of potential genetic markers for the diagnosis of such conditions.

**Figure 6.**
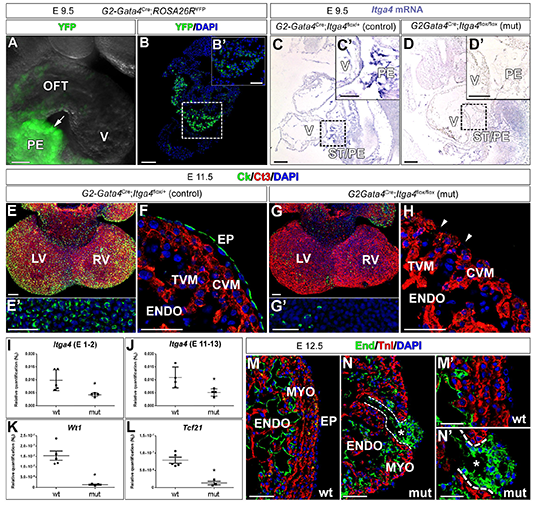
A developmental model for cells dynamics during CAF formation. **A, B**. Formation of the mouse epicardium in *Itga4* mutants (B) is disrupted as compared with the control condition (A). Loss of epicardial tissue reduces myocardial proliferation and results in ventricular myocardial wall discontinuities, local endocardial extrusion (B, arrow) and leads to embryonic death. **C-E**. In avian embryos, cryocauterisation-induced myocardial discontinuities precede epicardial formation and growth, which is originally normal. Blood pressure (D, red arrows) pushes and extrudes the endocardium towards the pericardial cavity, forming a pouch-like structure that resembles a single CAF (D, black arrow). Precocious endocardial-epicardial contact promotes the differentiation of smooth muscle and fibroblastic cells from epicardial tissues, which were originally fated to contribute to the coronary vascular system. Green, endocardium; blue, epicardium; red, myocardium.

## Acknowledgements

The authors thank Dr. A. Rojas (CABIMER, Sevilla, Spain) and Prof. Thalia Papayannopoulou (University of Washington, WA, USA) for sharing with us the *G2-Gata4-Cre* and *Itga4-floxed* mouse lines, respectively. We also thank Vanessa Benhamo (Institut Imagine) for her expert support with HREM. Finally, we thank all members of “DeCA” laboratory (University of Málaga, Málaga, Spain), and the “Heart Morphogenesis” laboratory (Institut Imagine and Institut Pasteur, Paris, France) for their help and fruitful discussions on this paper.

## Author contribution

JMPP designed the whole study and drafted the manuscript; PPG, BB, VB and LH carried out experiments; PPG, ARV, JAG, LC, AO, BP, DMG, JPR, DGC, SM, JLP and JMPP conceived experiments and analysed data. JMPP and JAGD provided funding. All authors were involved in writing the paper and had final approval of the submitted and published versions.

