## Supplementary figures and images for "Origin of congenital coronary arterio-ventricular fistulae from anomalous epicardial and myocardial development"

### Supplemental figure 1

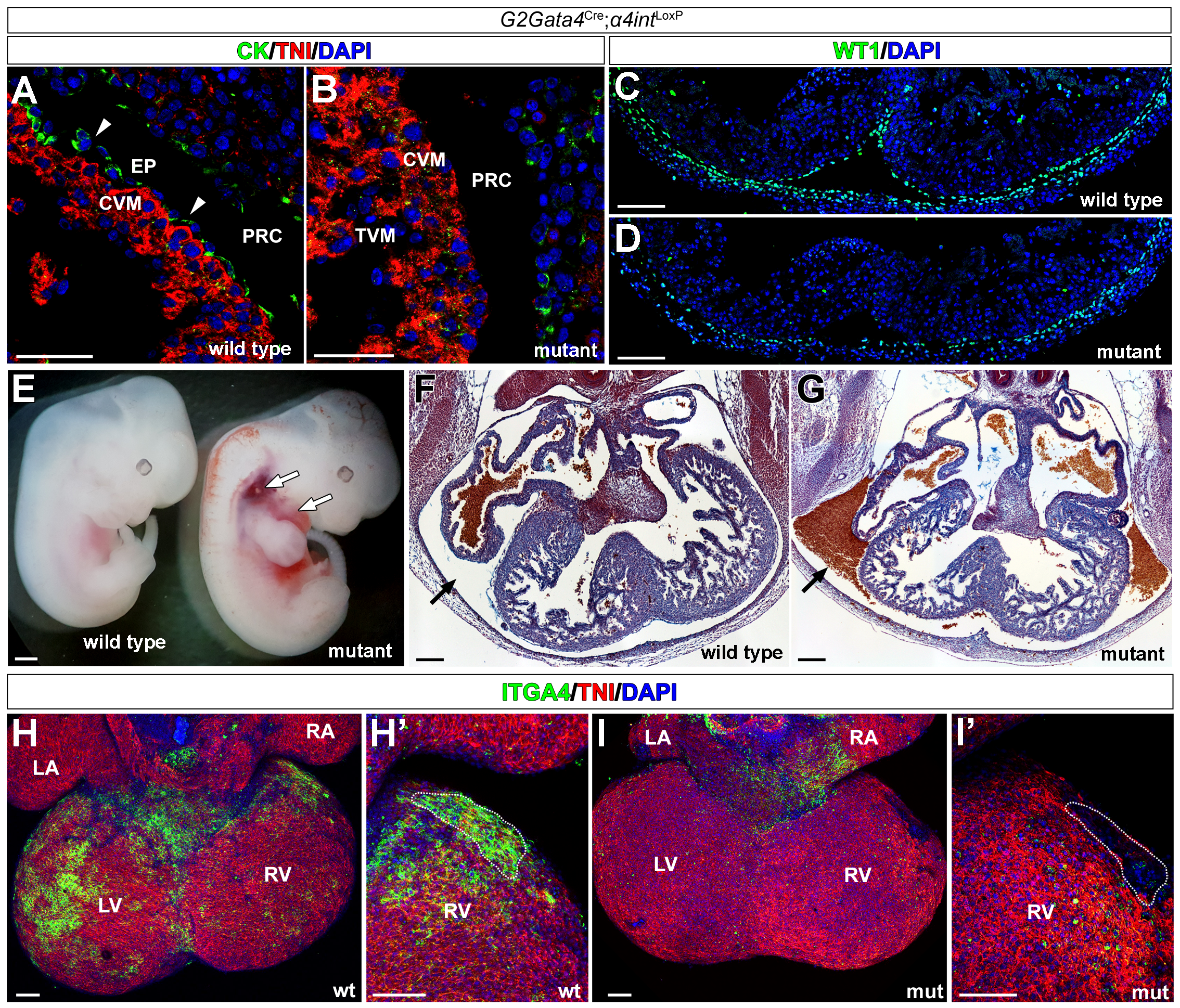

### Supplemental figure 2

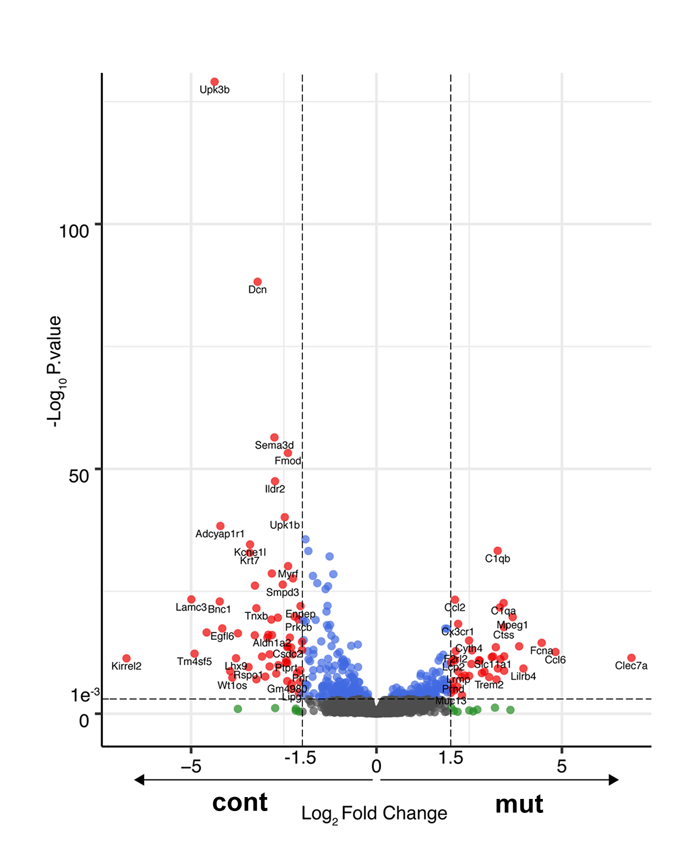

### Supplemental figure 3

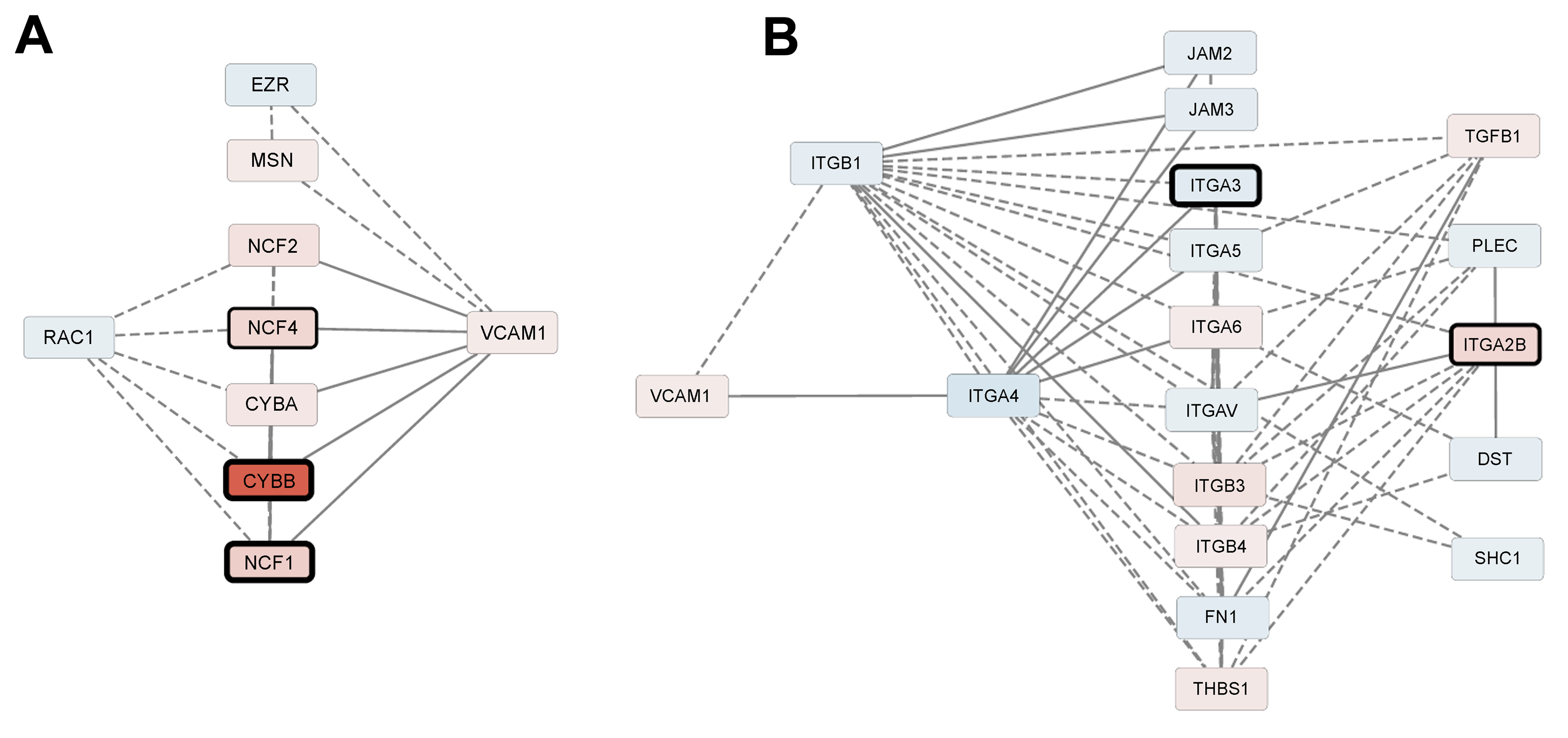

### Supplemental figure 4

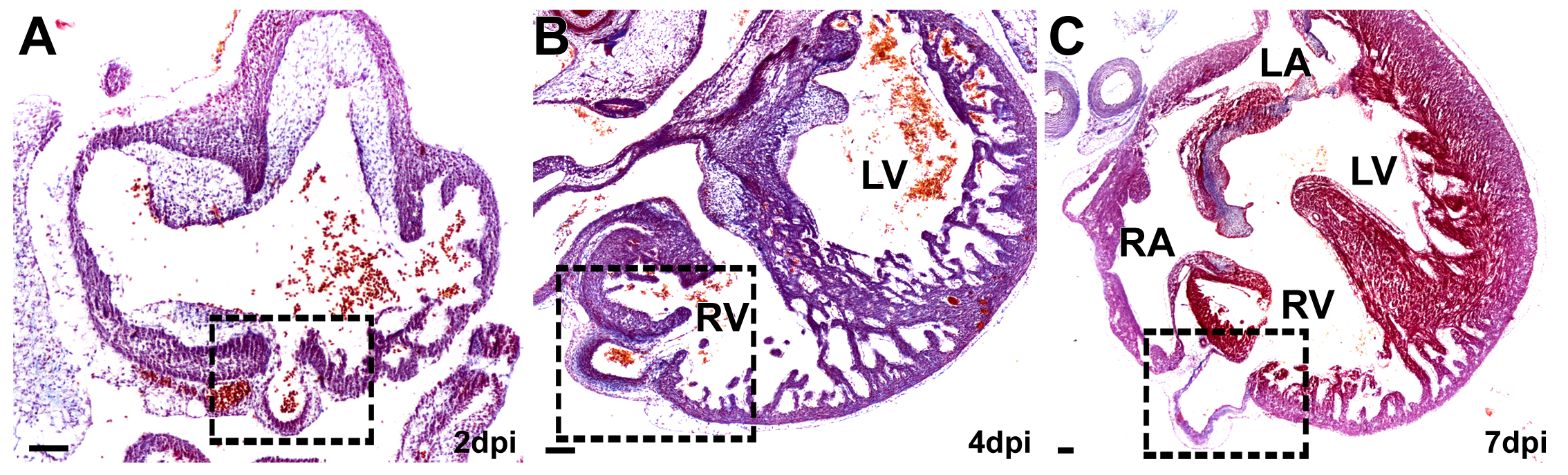
