## Supplementary information for "Origin of congenital coronary arterio-ventricular fistulae from anomalous epicardial and myocardial development"

**SUPPLEMENTARY INFORMATION (Palmquist-Gomes et al.)**

***SUPPLEMENTARY METHODS***

**In situ hybridization (ISH)**

E9.5 embryos were fixed in 4% paraformaldehyde/PBS, dehydrated in an ethanol series, paraffin-embedded, sectioned at 10 µm, and mounted on aminoalkylsilane-coated slides. Dewaxed sections were treated for proteolytic digestion with 20 µg/ml proteinase K in PBS (7 minutes at 37ºC), washed in 0.2 % glycine/PBS (5 minutes) and in PBS (two times, 5 minutes each). Sections were then post-fixed for 10 min in 4% formaldehyde/0.2% glutaraldehyde/PBS, washed twice in PBS for 5 min, and submitted to a pre-hybridization step in the hybridization mix [50% formamide, 5xSSC (saline-sodium citrate buffer), 1% block solution (Roche), 5 mM EDTA, 0.1% Tween-20, 0.1% Chaps (Sigma; St. Louis, MO), 0.1 mg/ml heparin (Becton-Dickinson; Mountain View, CA), and 1 mg/ml yeast total RNA (Roche)] without probe for 1 hour at 70ºC, and then hybridized with a probe against Itga4 (1ng/ml) overnight at 70ºC. After hybridization, the sections were washed in 2x SSC (pH7), washed twice in 50% formamide/2x SSC, pH7, 65ºC (30min and 1 hour, respectively) and three times in TNT (10 min each). Probe tissue binding was immunologically detected using a sheep anti-digoxigenin Fab covalently coupled to alkaline phosphatase and NBT/BCIP as chromogenic substrate, according to the manufacturer's protocol (Roche). Samples were incubated in NBT/BCIP 7 hours, washed and dehydrated using an ethanol series (50º, 70º, 80º, 90º and 96º, one minute each). Samples were finally washed in absolute ethanol (3x1'), an ethanol:xylene 1:1 solution (1') and 5' in absolute xylene, and mounted in EUKITT mounting medium.

**Trichrome staining**

Mallory's Trichrome staining includes the sequential incubation of the samples in corrosive sublimate solution (distilled water saturated with mercury chloride) for 30 minutes, in 1% acid fuchsine for 30 seconds, 1% phosphomolybdic acid for 75 seconds, and in Mallory's liquid (2.5% orange G, 2% oxalic acid and 0.5% aniline blue) for 45 seconds. All samples were gently washed with distilled water after each staining step. Stained tissue slides were washed in pure ethanol, xylene and mounted in DPX medium (BDH; Cat No. 361254D).

**Immunofluorescence analyses of mouse embryos**

For immunofluorescence analyses, non-specific binding sites were blocked in 16% goat serum, 1% bovine serum albumin, and 0.5% Triton X-100 in Tris-PBS (SBT) for 1 hour at room temperature (RT). Goat serum was replaced by horse serum in Vcam-1 immunostaining. Samples were incubated with primary antibodies (Table1) overnight at 4ºC. Samples were washed in PBS and incubated with secondary antibodies (Table 2) for 1 hour at room temperature. Nuclei were counterstained with DAPI (1/2000 dilution, Sigma D9542) and all samples analysed under a TCS-NT-laser confocal microscope (SP5, LEICA).

***BrdU tissue incorporation, immunohistochemistry and quantification***

Paraffin sections from BrdU injected embryos were treated with 2N HCl (30’ at room temperature) and washed in 100mM sodium tetraborate (5’). Samples were blocked with goat serum and incubated overnight in the corresponding primary antibodies (BrdU and troponin I; Table 1) at 4ºC. Samples were then washed in PBS and secondary antibodies (Table 2), and incubated for 1 hour at room temperature. All sections were mounted in a 1:1 TPBS-glycerol solution and analysed under a laser confocal microscope (SP5, LEICA). Proliferating cardiomyocytes were estimated with the IMARIS® software. First, the outer compact myocardium was separated from the embryonic ventricle (TNI+) for the analysis using Adobe Photoshop® software. All nuclei (DAPI+) and proliferating nuclei (DAPI+/BrdU+) in TNI+ cells (cardiomyocytes) were counted to calculate the compact myocardium proliferation rate. Three different sections of five different hearts (n=5) were analyzed in each condition. Final values of each replicate represents the mean of the values from the three sections measurements.

**qPCR protocol**

1 µg of total RNA was converted into cDNA using oligo dT Primer (2.5 µM) and Random primers (5 µM) following the manufacturing instructions (PrimeScriptTM

RT reagent Kit, Takara). For each qPCR the cDNA equivalent to 5 ng RNA was used. The qPCR reactions contained power SYBR green PCR master mix (Applied Biosystems) and an equimolar primer mix (0.8 μM). The amplification protocol consisted of 2 min at 50°C, 10 min at 95°C, followed by 40 cycles of 15 seconds at 95°C, and 1 min at 60°C, and completed with a standard melting curve protocol (15 seconds at 95°C, 1 min at 60°C and 15 seconds at 95°C).

**Culture of embryonic proepicardium and endocardium**

Embryonic quail hearts were isolated at HH16-17 and incubated at 37°C, 5% CO2 in a hanging drop (20μL) culture system using Dulbecco's modified eagle medium (DMEM)-High Glucose (GIBCO) supplemented with 10% heat-inactivated foetal bovine serum (FBS), 2% heat-inactivated chick serum and 100 units/ml penicillin/streptomycin DMEM, 1% glutamine, 10% FBS, 2% chick serum and 1% penicillin/streptomycin, as culture medium. Some of the quail ventricles were digested with trypsin (5 min. at 37ºC), centrifuged (5 min. at 1500rpm), washed in DMEM and cultured at 37°C, 5% CO2 in a hanging drop (20μL) for three days until cardiac tissue aggregates were formed. Co-cultures were set up culturing together freshly excised chick proepicardia and quail ventricles (or cardiac aggregates) in hanging drops (20μL) using the culture medium described above.

**Immunochemical analyses of the human paediatric CAF**

For single immunoperoxidase staining of these human tissues, endogenous peroxidase activity was quenched by incubating the sections for 30 minutes in 3% hydrogen peroxide. After washing, endogenous biotin was blocked using a specific avidin–biotin blocking kit (Vector SP2001). Non-specific binding sites were saturated for 1 hour with 16% sheep serum, 1% bovine serum albumin, and 0.5% Triton X-100 in Tris-PBS (SBT) at room temperature. Slides were then incubated overnight at 4°C in anti-Von Willebrand factor primary antibody (Table 1), washed in PBS, incubated for 1 hour at room temperature in biotin-conjugated goat anti-rabbit IgG (Table 2) and washed again in PBS. After a final incubation (1 hour, room temperature) in streptavidin-peroxidase complex (SIGMA S5512), sections were washed, and peroxidase activity was developed using SIGMAFASTTM 3,3'-diaminobenzidine tablets (SIGMA D4293). Tissues were counterstained with Harris haematoxylin. For αSMA immunofluorescence, non-specific binding sites were blocked with SBT and slides were incubated overnight in the primary antibody (Table 1) at 4°C. Samples were washed in PBS and incubated for 1 hour at room temperature in the secondary antibody (Table 2; goat anti-mouse FITC). All cell nuclei were counterstained using DAPI (1/2000, Sigma D9542), tissue sections were mounted in a 1:1 glycerol/TPBS solution and analysed under a TCS-NT laser confocal microscope (SP5, Leica).

***Antibodies***

**Supplementary table 1.** Primary antibodies used in this work.

| Epitope | host | clonality | dilution | reference |
|---|---|---|---|---|
| α-smooth muscle actin | mouse | monoclonal | 1/200 | SIGMA A2547 |
| BrdU | mouse | monoclonal | 1/100 | DSHB G3G4 |
| Cytokeratin | rabbit | polyclonal | 1/200 | DAKO Z0622 |
| Endomucin | rat | polyclonal | 1/500 | Santa Cruz 65495 |
| QCPN | mouse | monoclonal | 1/20 | DSHB AB_531886 |
| QH1 | mouse | monoclonal | 1/100 | DSHB AB_531829 |
| Troponin I | rabbit | polyclonal | 1/100 | Santa Cruz 15368 |
| Troponin I | mouse | monoclonal | 1/100 | Santa Cruz 365446 |
| Troponin T | mouse | monoclonal | 1/100 | DSHB CT3 |
| Vcam1/CD106 | goat | polyclonal | 1/200 | R&D systems AF643 |
| Von Willebrand factor | rabbit | polyclonal | 1/800 | SIGMA F3520 |

**Supplementary table 2.** Secondary antibodies used in this work.

| Epitope | host | Conjugated molecule | dilution | reference |
|---|---|---|---|---|
| Mouse IgG | goat | FITC | 1/200 | SIGMA F2012 |
| Mouse IgG | donkey | Alexa Fluor® 647 | 1/200 | Jackson IR 715-605-151 |

| | | | | |
|---|---|---|---|---|
| Rat IgG | donkey | Alexa Fluor® 488 | 1/200 | Jackson IR 712-545-153 |
| Rabbit IgG | goat | Alexa Fluor® 647 | 1/200 | Jackson IR 711-605-152 |
| Rabbit IgG | goat | biotin | 1/200 | SIGMA B7389 |

### *PCR*

**Supplementary table 3.** Primer sequences for qPCR analyses. **Abbreviations**: Amp, amplicon size; Temp, template.

| Name | Temp | FW | RV | Amp |
|---|---|---|---|---|
| *α4 integrin* (E1-2) | NM_010576.4 | GTTGTACTTCGGGGTGCCAA | CCAGGATTGACCACTGAG | 189 |
| *α4 integrin* (E12-13) | NM_010576.4 | AATCTCCTCCACCTACTCACAG | GCACCAACGGCTACATCAAC | 129 |
| *Tcf21* | NM_011545.2 | ACAAGTACGAGAACGGTTACATT | CAGGTCATTCTCTGGTTTG | 80 |
| *Vcam1* | NM_011693.3 | CAAAAAGGGACGATTCCG | GGCACATTTCCACAAGTG | 183 |
| *Wt1* | NM_144783.2 | ATACCAAATGACCTCCCAGC | GCCACTCCAGATACACGC | 243 |

### *SUPPLEMENTARY FIGURES*

**Supplementary figure 1. *Itga4* ablation in *G2-Gata4*$^{+}$ cells disrupts epicardial and myocardial embryonic development.** E10.5 epicardial cells are Ck$^{+}$ (A, arrowheads). Ck$^{+}$ epicardial cells are scarce in stage-matched *G2-Gata4*$^{Cre/+}$;*Itga4*$^{flox/flox}$ mutants, but no myocardial abnormalities were observed in these animals (B). Wt1 protein expression is evident in the developing epicardium of E11.5 *G2-Gata4*$^{+/+}$;*Itga4*$^{flox/+}$ control embryos (C), while the number of Wt1$^{+}$ epicardial cells in E11.5 mutant embryos is significantly reduced (D). Pericardial hemorrhage is evident in all analysed E12.5 mutants, both macroscopically (E) and after histological inspection (F,G, arrows). *Itga4* epicardial deletion results in a marked decrease of α4-integrin protein in E12.5 mutant embryonic epicardium (compare H with I). The absence of α4-integrin$^{+}$ epicardial cell clusters correlates with the presence of myocardial discontinuities in mutant ventricles (compare H' with I'). **Abbreviations:** Ck, cytokeratin; CVM, compact ventricular myocardium;

DAPI, diamidino-2-phenylindole; END, endocardium; EP, epicardium; LV, left ventricle; PRC, pericardial cavity; RV, right ventricle; TnI, troponin I; TVM, trabeculated ventricular myocardium; Wt1, Wilms' tumor protein 1. **Scale bars:** A,B: 50µm; C,D,F,G,H,H',I,I': 100µm; E: 500µm.

**Supplementary figure 2.** RNA-seq analysis was performed on *G2-Gata4*$^{\mathbf{Cre/+}}$;*Itga4*$^{\mathbf{flox/flox}}$ (mutant; n=4) and *G2-Gata4*$^{+/+}$;*Itga4*$^{\mathbf{flox/+}}$ (control; n=5) E11.5 ventricles (each replicate represents a pool of three ventricles). All differentially expressed genes (DEG) in control vs mutant ventricles are represented in a volcano plot.

**Supplementary figure 3. Protein interaction models involving VCAM1 and ITGA4.** A protein-protein interaction (PPI) model was generated using proteins whose mRNA profile was available in our RNA-seq data (B,C). This model was downloaded from StringDB and visualized in Cytospace. Nodes were selected based on first and second PPI associations with VCAM1 (A) and ITGA4 (B), using a maximum of 10 connections derived from "experiments" or "databases". Red and blue colors denote gene overexpression and underexpression in our RNAseq analysis, respectively. Bold box borders denote differential expression based on adjusted p-value < 0.05. Lines highlight direct protein interaction; dashed lines denote protein interactions as based on experimental evidence, while solid lines result from database evidence. Nodes have been positioned to highlight VCAM1 (B) and ITGA4 interactions (C). VCAM1 is also included in panel C due to its relation with ITGB1.

**Supplementary figure 4. Cryoinjury of the quail embryonic heart leads to the formation of CAF-like structures.** Mallory's trichrome staining of quail embryonic hearts. **A-D**. Transverse sections of cryoinjured quail embryonic hearts at 2-, 4- and 7-days post injury (dpi) show fistula-like structures on their ventricular walls (A-C; boxed area). **Abbreviations:** dpi, days post injury; LA, left atrium; LV, left ventricle; RA, right atrium; RV, right ventricle. **Scale bars:** 100µm.
